# Engineering Cybergenetic Cell-Based Therapies

**DOI:** 10.64898/2026.01.16.699922

**Authors:** Ching-Hsiang Chang, Asterios Arampatzis, Samuel Balula, Mucun Hou, Maurice Filo, Mingzhe Chen, Federica Cella, Mustafa Khammash

## Abstract

Adaptive, closed-loop control of cellular behavior is essential for next-generation therapies, yet most current treatments operate in an open-loop manner and lack robustness to patient variability and disease dynamics. Here, we establish a control-theoretic platform for rational engineering of closed-loop cell-based therapies that achieve precise and robust regulation. First, we introduce multi-dimensional nullgram profiling, a high-throughput approach that enables quantitative prediction and design of advanced genetic controllers in human cells across circuit topologies and parameter regimes in a single experiment. To evaluate dynamic therapeutic behavior, we next develop Cyberpatient-in-the-loop, an optogenetic digital twin platform that interfaces engineered mammalian cells with computational disease models, enabling systematic testing of closed-loop performance under realistic perturbations. Finally, we leverage these approaches to implement integral feedback cell therapies that sense inflammatory signals and autonomously regulate cytokine levels in primary immune cell cultures. Together, these results establish a general paradigm for engineering cell-based control systems and provide a foundation for next-generation cell therapies.

---

Homeostasis, the ability of biological systems to maintain internal stability in the presence of external perturbations, is fundamental to health (*1–3*). Disruption of homeostatic regulation can lead to chronic inflammation, metabolic dysfunction, and neurodegenerative disease, often necessitating long-term therapeutic intervention (*4–6*). However, most existing treatments do not actively restore regulatory feedback. Instead, therapeutics are administered in an open-loop manner, without continuous sensing of disease state or dynamic adjustment of dosage. As a result, achieving precise regulation while maintaining robustness to patient variability and disease state fluctuations remains a major clinical challenge, particularly for drugs with narrow therapeutic windows and high interpatient variability (*7*).

An appealing alternative is the development of *cybergenetic* cellular therapies capable of autonomous, closed-loop regulation (*8*). In this paradigm (Fig. 1A), living cells are engineered as biomolecular control systems that (i) sense disease-relevant biomarkers, (ii) process these signals in real-time through embedded molecular control circuits, and (iii) dynamically modulate therapeutic outputs to maintain a desired physiological setpoint. By embedding feedback directly at the cellular level, cybergenetic cells have the potential to reconstitute disrupted homeostatic loops rather than merely counteracting downstream symptoms. Advances in synthetic biology have made this vision increasingly feasible (*9–13*), as illustrated by recent clinical successes of engineered cell-based therapies (*14, 15*). Nonetheless, translating closed-loop control principles into reliable and application-ready cell therapies remains challenging, as achieving both precise setpoint regulation and robust performance across operating regimes requires careful control system design (*16*).

**Figure 1:**
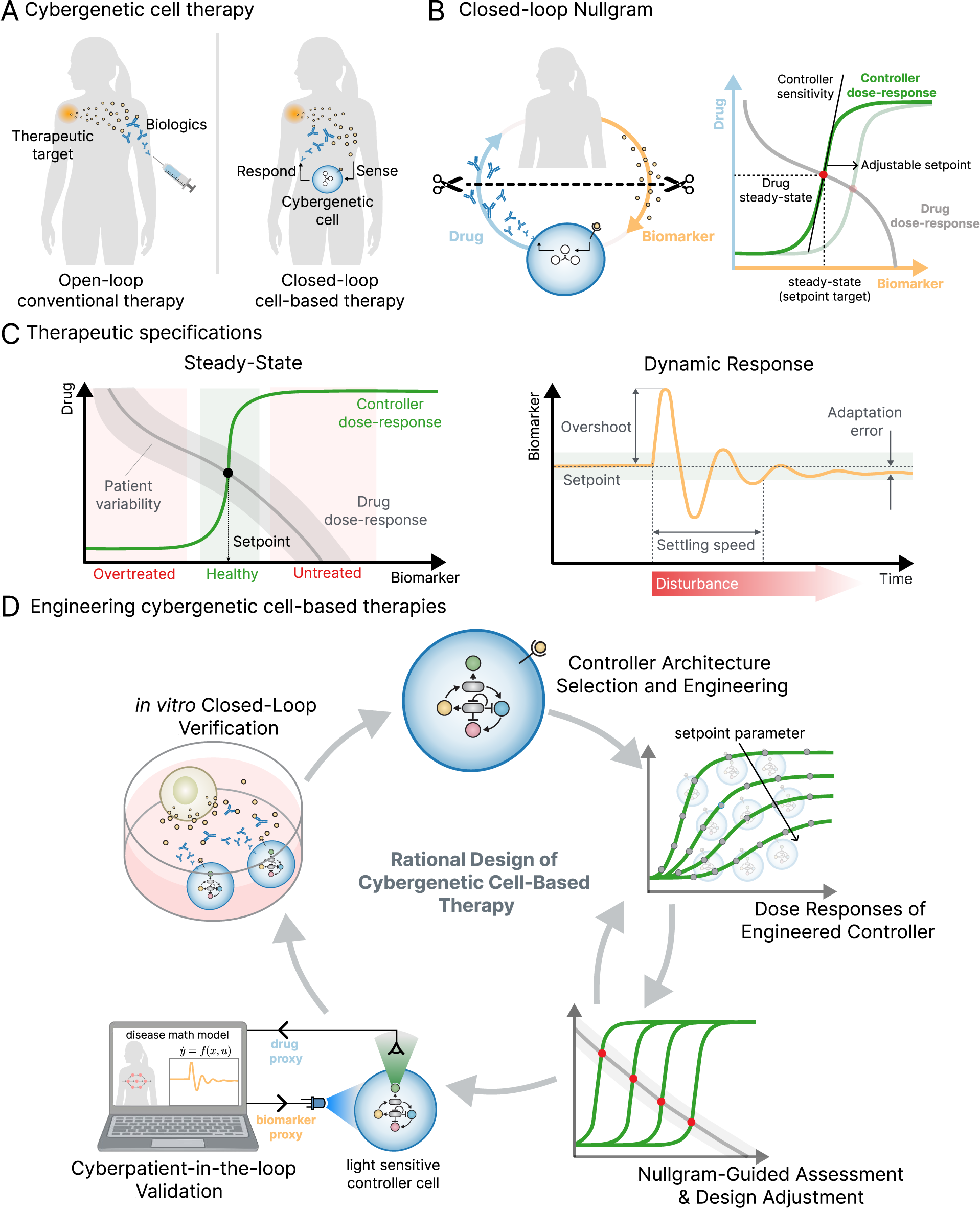
A control-theoretic framework for predictive, robust, and precise design of closedloop cell-based therapies. (A) Conceptual comparison between conventional open-loop molecular therapy and closed-loop cell-based therapy. In conventional treatment, biologics are administered without feedback from the disease state, leading to sensitivity to patient variability. In closed-loop therapy, engineered cybergenetic cells sense a disease biomarker, compute a control action, and autonomously regulate therapeutic output to maintain homeostasis. (B) Closed-loop nullgram framework. Controller dose–response and process (drug) dose–response curves are overlaid in biomarker–drug space. Their intersection defines the closed-loop steady state, which can be predictively tuned through controller setpoint and sensitivity. (C) Control specifications for therapy design. Left: Steady-state specification, in which the controller dose–response must compensate for patient variability and place the closed-loop equilibrium (setpoint) within a sensitive operating regime. Right: Dynamic specification, in which closed-loop responses must reject disturbances while ensuring stability, fast settling, limited overshoot, and robust perfect adaptation. (D) Rational design workflow for cybergenetic cell-based therapy developed in this work.

Negative feedback motifs have long been employed in synthetic biology to synthetically realize homeostasis (*17–19*). Among these, integral feedback motifs are particularly attractive because they enable systems to robustly return to predefined setpoints despite persistent disturbances, thereby achieving Robust Perfect Adaptation (RPA) (*20*). One minimalist yet powerful implementation is the Antithetic Integral Controller (AIC) (*21–23*). It uses two molecular species that mutually sequester, with one setting a reference signal and the other sensing the regulated output. This topology has been realized in cell-free systems, bacterial and mammalian cells (*22, 24–29*).

Integral feedback controllers that enable precise and robust regulation are highly attractive for therapeutic applications. One biological signal that highlights the need for such control is tumor necrosis factor–*α* (TNF-*α*), a central pro-inflammatory cytokine whose concentration must be tightly confined within a narrow physiological range (*30*). Excess TNF-*α* drives chronic inflammation and autoimmunity, whereas insufficient levels impair host defense (*31, 32*). We engineered genomically integrated cell lines with our previously developed intein-based AIC (*26*), coupling an NF-κB–responsive promoter as the sensor, and adalimumab, an anti–TNF-*α* biologic, as the actuator (Supplementary Fig. S1). The key reaction underlying the controller is intein splicing, which implements the antithetic motif and concurrently generates a DNA-binding splice product that introduces additional repression. Together, these features result in a biological proportional–in-tegral–derivative (PID) controller, as confirmed by theoretical analysis (Linear Perturbation Analysis in Supplementary Text). PID controllers promise faster, more stable, and finely tunable responses (*11, 33, 34*), motivating us to evaluate our controller’s ability to regulate TNF-*α* in co-culture assays (Supplementary Fig. S1D). While the circuit exhibited broad setpoint tunability, robust TNF-*α* homeostasis was only achieved at low setpoints, suggesting performance breakdown at higher setpoint regimes.

This divergence reflects a common translational challenge. Controllers that perform well in simple, idealized systems often operate within narrower ranges in physiologically relevant contexts, where dilution, saturation, and other non-ideal effects dominate (*24, 35–38*). In this setting, the rational design and tuning of more advanced architectures, such as PID controllers, become particularly difficult. While these controllers promise improved performance, they also introduce greater implementation complexity and rely heavily on precise parameter tuning (*34*). Without tools to dissect controller behavior and guide tuning, their advantages are difficult to realize in complex biological settings. To overcome these limitations, systematic strategies are needed to evaluate and predict controller performance across designs and operating regimes, thereby bypassing the slow design–build–test–learn cycle and accelerating translation.

Here, we develop an engineering framework for the rational and scalable design of homeostatic cell therapies, inspired by principles from control theory and synthetic biology. The strategy is summarized in Figure 1, highlighting the contrast between open-and closed-loop therapies, the steady-state and dynamic performance specifications required for therapeutic control, and the integrated design and validation workflow developed in this work. Guided by the mechanistic insights obtained, we engineer and deploy novel closed-loop cell therapies for immune system regulation.

## 1 Multi-dimensional nullgram profiling predicts steady-state controller behavior

In closed-loop biological systems, the observed behavior does not arise from the controller or the target process in isolation, but from their mutual interaction once coupled through feedback. A central challenge in closed-loop cell therapy design is to predict, prior to full circuit construction, whether a given controller architecture and parameter regime can achieve precise and robust regulation once embedded in feedback.

The steady-state behavior of coupled dynamical systems, including feedback control systems, has been studied extensively (*8, 39–41*). Fixed points of such systems can be characterized graphically as the intersections of their nullclines, a classical insight that has been successfully exploited in the experimental analysis and design of synthetic biological circuits (*42–44*). Here, we introduce a multidimensional nullgram framework for mammalian feedback controllers, providing an experimentally accessible input–output representation that exploits natural variability to enable steadystate analysis of entire families of controllers within a single experiment.

Let *y* = *ϕ*(*u*) denote the steady-state input–output map of the control circuit, experimentally measured as a steady-state dose–response curve. Similarly, let *u* = *π* (*y*) denote the steady-state input–output map of the target process to be controlled. Conceptually inspired by nullcline analysis (*39*), we define a nullgram to be the experimentally derived steady-state map obtained by plotting both relationships, *ϕ*(·) and *π* (·), on the same (*u, y*) axes, such that their intersection directly predicts the closed-loop equilibrium (Fig. 1B). Nullgrams reveal how regulatory components balance at steady state: the controller dose–response curve characterizes how actuator (e.g. drug) production varies with biomarker levels, while the process dose–response curve describes how the regulated target responds to actuator activity. Their intersection defines the closed-loop equilibrium, which must lie within a sensitive region of the controller curve to ensure robust regulation.

Building on this definition, we extend nullgram profiling to a multidimensional setting by profiling entire families of controllers against a shared target process (as summarized in Box 1). In this formulation, natural cell-to-cell variability and tunable controller parameters jointly span a multidimensional steady-state space, allowing the input–output properties of multiple controller designs to be experimentally separated and recombined. This multidimensional nullgram profiling yields a mathematically precise yet experimentally accessible framework for predicting closed-loop performance across controller architectures and operating regimes.

To validate the predictive accuracy of nullgram profiling, we first defined a minimal closed-loop system that is both biologically realizable and analytically interpretable, which served as an experimental ground truth. In this system, the controller regulates mRNA expression through an intein-based activator–splicer architecture. The controller receives three independent signals: a constitutive reference defining the setpoint, the mRNA level serving as the controller input, and a tunable gain that scales controller strength. The controller compares the reference with the controller input through intein-mediated splicing and modulates mRNA expression accordingly, with the magnitude of the response determined by the controller gain. Each of these three signals was independently observable through fluorescent protein proxies, enabling direct experimental readout of the setpoint (RFP), gain (BFP), and controller input (FRFP) (Fig. 2A).

**Figure 2:**
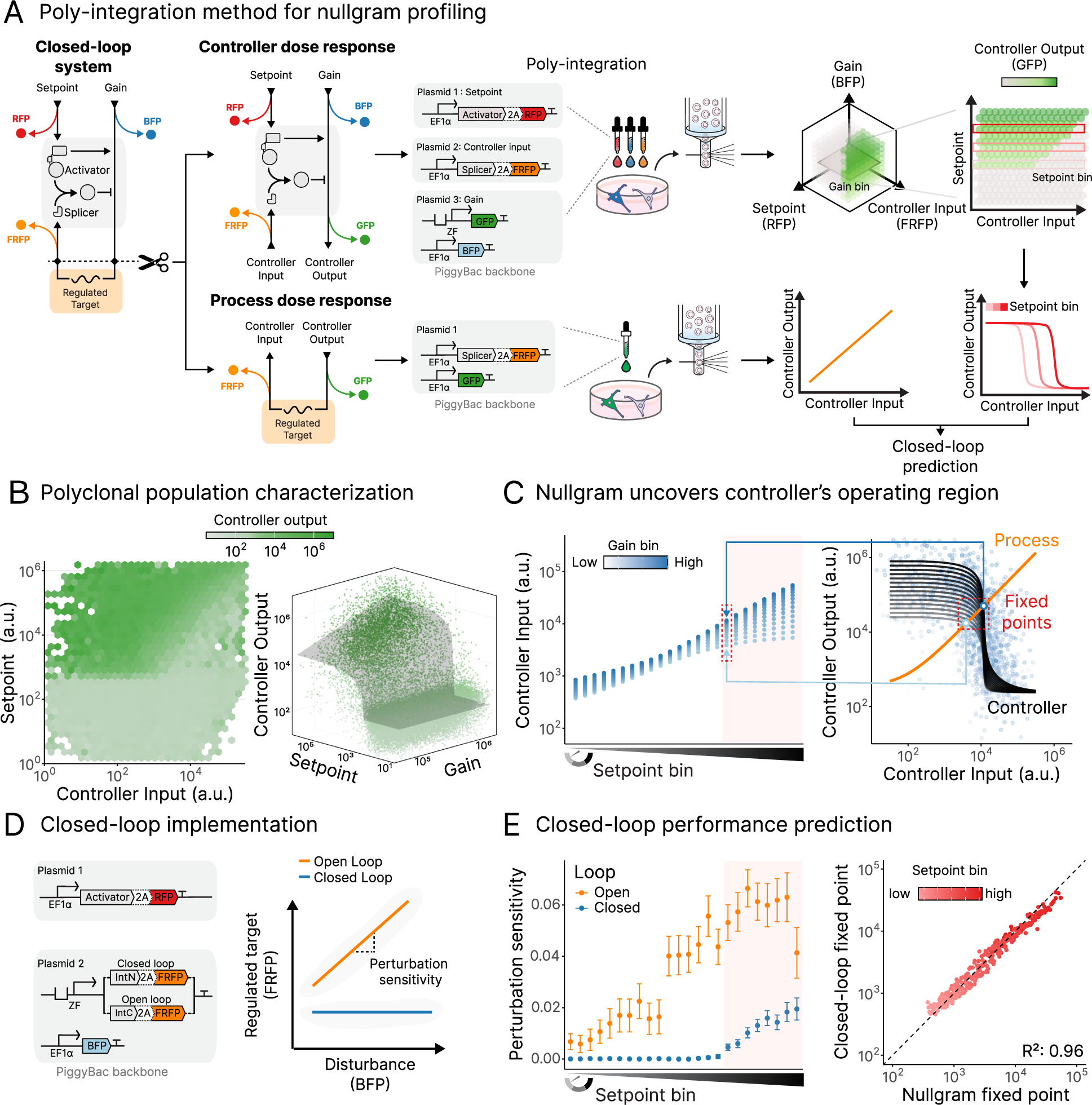
Multi-dimensional nullgram profiling predicts closed-loop behavior of genetic controllers. (A) Construction of a nullgram using poly-integration. A minimal intein-based closedloop system is decomposed into a controller and a process, whose steady-state dose–response relationships are independently measured via PiggyBac-mediated poly-integration. The controller integrates three inputs—a constitutive setpoint reference (RFP), the controller input (FRFP), and a tunable gain (BFP)—and produces an output reported by GFP. The process dose–response curve maps controller output to the regulated target. Together, these relations define the nullgram and predict closed-loop equilibria. (B) Variability-enabled reconstruction of controller dose–response surfaces. Poly-integration generates polyclonal populations spanning wide ranges of setpoint, input, and gain, populating a three-dimensional input–output space at single-cell resolution. Left: Projection onto the input–setpoint plane reveals a sharp diagonal boundary consistent with intein-mediated splicing; hexagons denote binned mean output levels. Right: A three-input, one-output mechanistic model yields smooth controller dose–response surfaces; points represent individual cells and the gray surface indicates the fit. (C) Identification of controller operating regimes. Right: Slicing the dataset at fixed setpoints yields families of controller dose–response curves across gains; overlay with the process curve defines a unique fixed point for each parameter combination. Left: Across setpoints, groups of fixed points corresponding to different gains are revealed, with gain indicated by color. (D) Closed-loop implementation. Left: The minimal closed-loop circuit is assembled from two donor plasmids encoding the reference and feedback modules; an open-loop analog is generated using a non-splicing intein. Right: In open-loop, regulated target levels increased monotonically with gain, whereas closed-loop circuits compensate for gain-induced perturbations, demonstrating disturbance rejection. Perturbation sensitivity is quantified as the slope of target output versus disturbance. (E) Quantitative prediction of steady-state performance. Left: Closed-loop control strongly reduces perturbation sensitivity, which rises sharply beyond a gain-dependent failure threshold at high setpoints. Points denote perturbation sensitivities estimated from linear fits (illustrated in Supplementary Fig. S2E), with 95% confidence intervals. Right: Nullgram-derived fixed points accurately predict closed-loop steady-state target levels across setpoints and gains. Each point represents a unique setpoint–gain bin, plotted as mean closed-loop output (y axis) versus the corresponding nullgram-predicted fixed point (x axis).

To extract the controller dose–response curve, we separated the controller from the process and characterized it in an open-loop configuration. In this setting, the controller retained its three inputs—setpoint, controller input, and gain—and produced controller output reported by GFP. To map controller input–output relationships across multi-dimensional parameter space, we developed a poly-integration strategy. Building on poly-transfection approaches (*45*) and PiggyBac-mediated genomic integration (*46*), poly-integration generates polyclonal cell populations spanning broad combinations of expression levels, thereby allowing reliable steady-state sampling across multidimensional input spaces (as summarized in Box 1).

#### Box 1. Primer on multi-dimensional controller dose-response profiling

Variability-enabled multidimensional dose–response profiling allows for high-throughput nullgram generation. Specifically, the controller under study receives three inputs: the control input *u* (orange), the setpoint *µ* (red), and the output gain *κ* (blue). In the experimental implementation, variability in DNA copy number at the loci encoding *µ* and *κ* (indicated by DNA icons) is modeled by random variables *N_µ_* and *N_k_*. Corresponding fluorescent reporters (*0*, *R*, *B*) allow the realized values of these variables to be measured in single cells. Variation in *N_u_* generates a sweep of the input *u* and thereby samples the controller’s steady-state input–output relation, while variation in *N_µ_*and *N_k_* provides conditioning on setpoint and gain. The controller output is given by *y* = *κ* · *f* (*Z*), reported by a green fluorescent protein (*G*); experimentally, this corresponds to the steady-state protein concentration, which reflects the underlying production rate scaled by dilution/degradation. Together, these steady-state dose–response curves define the controller input–output relation (the controller dose-response), parameterized by the setpoint *µ* and gain *κ*. By exploiting DNA copy-number variability, this approach enables reconstruction of this input–output relation across multiple parameter regimes in a single open-loop experiment, providing a basis for predicting closed-loop behavior once the controller is re-embedded in feedback.

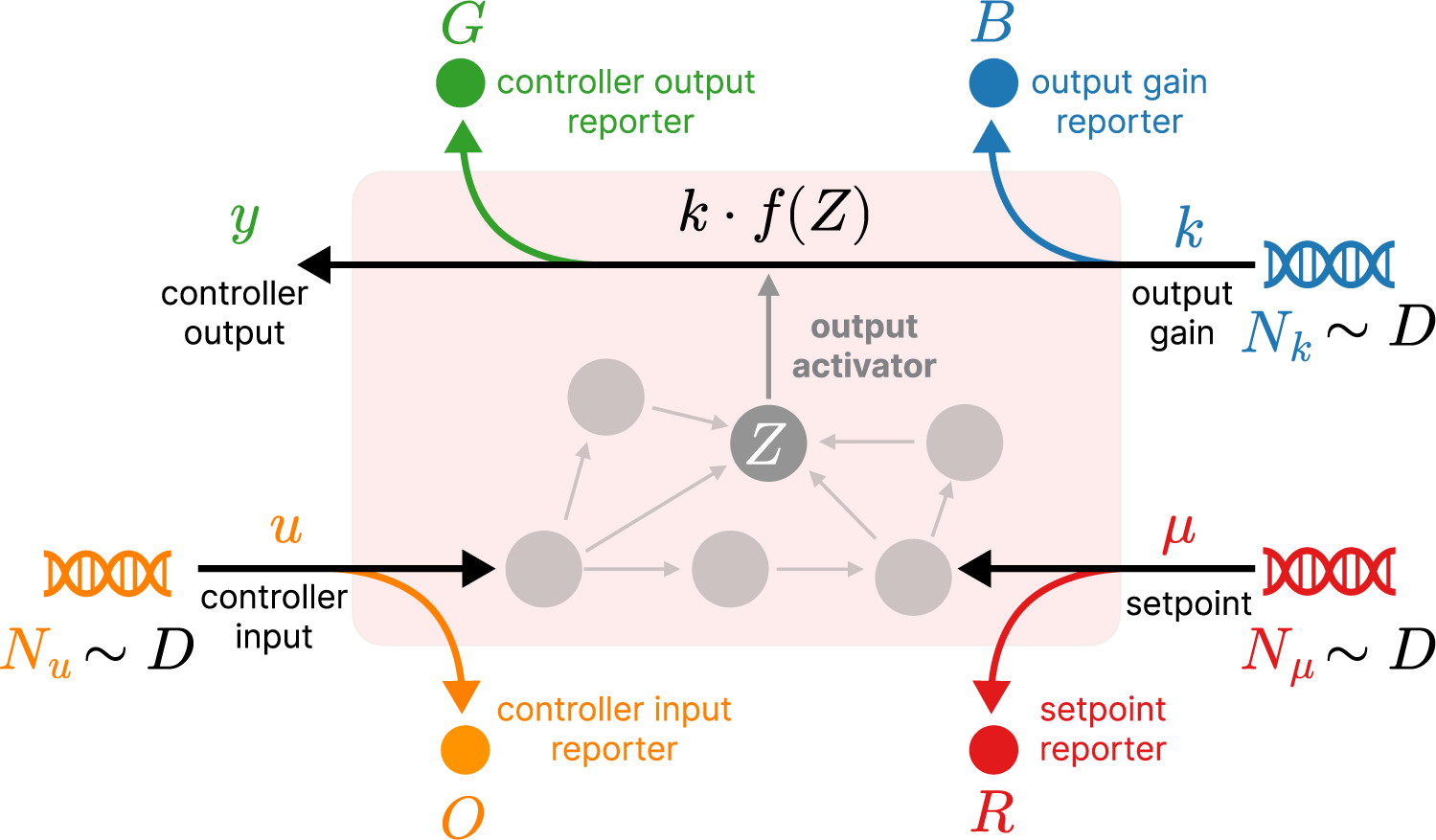

The experiment generates a three-dimensional scatter plot of single-cell measurements, with axes corresponding to the reporters *0* (input, orange), *R* (setpoint, red), and *B* (gain, blue). Each point represents an individual cell, with green intensity encoding the controller output reporter *G*. Slicing the dataset at a fixed gain value (for example, *B* = *B*_1_) selects cells with comparable controller strength. Projection of this slice onto the *0* − *R* plane reveals how the controller output varies jointly with input and setpoint. The resulting open-loop dose–response curves *G* (*0*), conditioned on selected setpoint values (for example, three representative setpoints indicated by horizontal red lines), exhibit characteristic sigmoidal shapes and systematic shifts with *R*. Together, this approach illustrates how DNA copy-number variability enables multidimensional reconstruction of controller dose–response curves across different setpoints and gains within a single experiment.

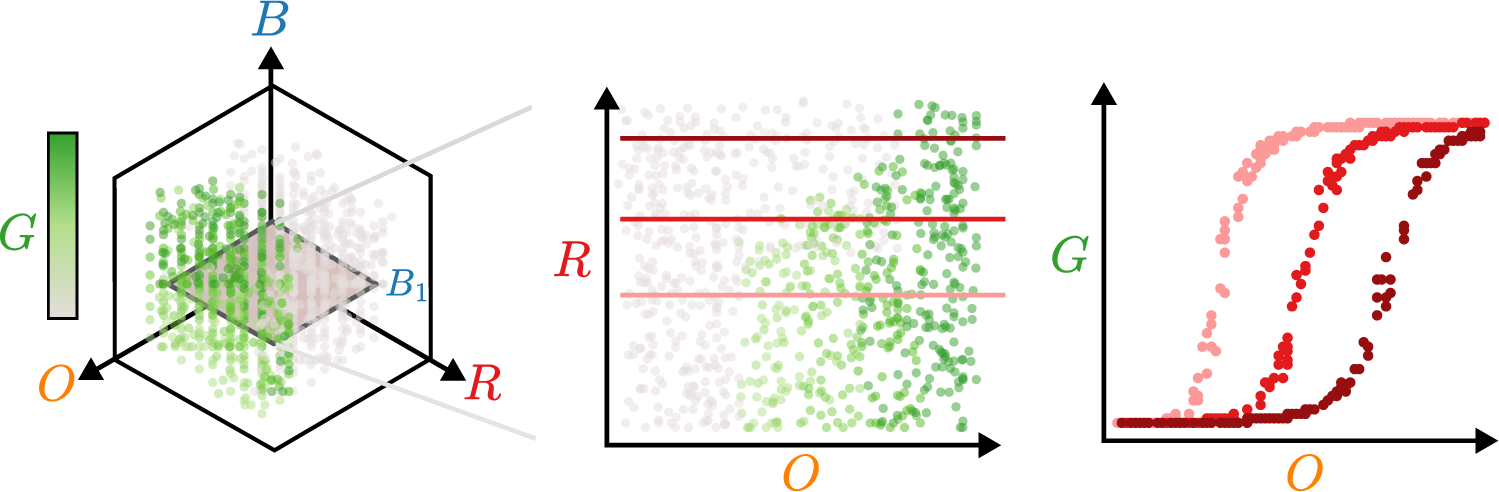

Three plasmids encoding corresponding inputs were randomly integrated using poly-integration, generating polyclonal populations spanning a wide range of input levels and producing diverse steady-state controller outputs (Supplementary Fig. S2A). Projection of single-cell data onto the input–setpoint plane revealed a sharp diagonal boundary in output expression, consistent with intein-mediated splicing (Fig. 2B, left). To derive smooth controller dose–response curves at defined setpoint and gain levels, we fit the entire dataset to a mechanistic three-input/one-output model (Fig. 2B, right; Supplementary Text).

In parallel, the process dose–response curve was constructed as a steady-state mapping between controller output (GFP) and controller input (FRFP) using a plasmid containing two constitutive units (Supplementary Fig. S2D). Overlaying the two dose–response curves yielded a single fixed point, whose steady-state level increased with the controller setpoint. High-setpoint controllers required higher gain to maintain the fixed point within the ultrasensitive operating regime; otherwise, controller saturation resulted in steady-state error (Fig. 2C).

To experimentally validate nullgram predictions, we constructed the minimal closed-loop circuit described above using two donor plasmids (Fig. 2D). One plasmid constitutively expressed the activator together with an RFP reporter, thereby defining the controller setpoint. The second plasmid implemented the feedback module, with an activator-responsive promoter driving IntN to close the loop and FRFP reporting the regulated target mRNA (controller input). Feedback plasmid copy number, serving as controller gain, was reported by an additional constitutive BFP expression unit. An open-loop analog was generated by replacing IntN with IntC, which cannot splice the activator. In open-loop, target levels increased with gain owing to the absence of feedback (Supplementary Fig. S2E). In contrast, closed-loop circuits compensated by producing additional IntN to suppress the activator, flattening the gain-induced disturbance–response curve. As predicted, high-setpoint controllers exhibited incomplete disturbance rejection when controller gain was insufficient to maintain the fixed point within the ultrasensitive operating regime (Fig. 2E, left and Supplementary Fig. S2E). Beyond qualitative agreement, nullgram profiling quantitatively predicted steadystate target levels for each combination of setpoint and gain with high accuracy (Fig. 2E, right). Together, these results show that nullgram profiling can experimentally derive the steady-state properties of control circuits and predict their operational regimes without requiring full closed-loop construction, offering a generalizable tool for rational controller design.

## 2 A digital twin platform allows for testing genetic control systems in disease settings

Nullgram profiling provides a predictive approach for designing and interpreting the steady-state behavior of synthetic biological controllers. However, therapeutic efficacy also depends critically on dynamic performance, including disturbance rejection, transient overshoot, and stability in the presence of disease-driven fluctuations (Fig. 1C). Motivated by approaches in digital twin modeling and pharmacokinetic/pharmacodynamic systems that enable quantitative descriptions of disease processes, we sought to extend nullgram-guided design into clinically relevant regimes. We developed *Cyberpatient-in-the-loop*, a hybrid *in silico*/*in vitro* platform that couples engineered mammalian cells to real-time disease models (Fig. 3A). Building on the Cyberloop framework (*47*), Cyberpatient-in-the-loop enables closed-loop interaction between living cells and computational models, allowing systematic evaluation of controller dynamics under realistic pharmacokinetic and pharmacodynamic constraints. These dynamic performance metrics are essential for therapeutic applications, where transient failures, such as excessive cytokine excursions, can have severe clinical consequences.

**Figure 3:**
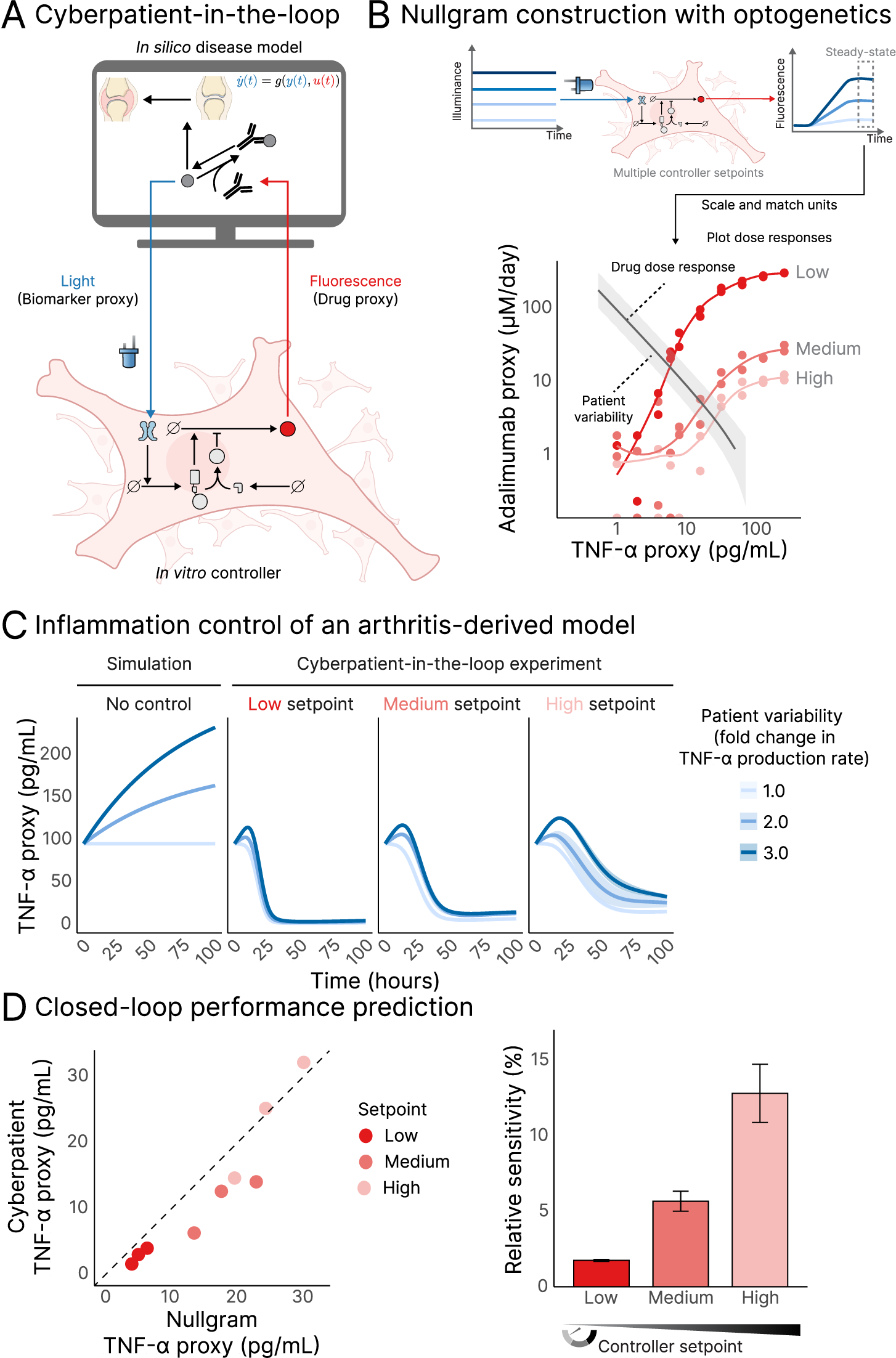
A digital twin platform enables predictive testing of genetic controllers in disease settings. (A) Schematic of the Cyberpatient-in-the-loop platform, coupling engineered optogenetic controller cell lines with an *in silico* disease model of interest. (B) Experimentally derived controller dose–response curves overlaid with the process dose–response curve, aligning cell-based dose–response with simulated pharmacodynamics. Points represent experimental steady-state data for two biological replicates and colored lines are smooth fits. The grey shaded region is derived from the pharmacodynamic model component of (*48*), with upper bound: simulation with TNF-*α* production rate *k_s_* = 6 · *k_s_* and lower bound: simulation with *k_s_*. Both axes are on log scales. (C) Closed-loop inflammation control, comparing open-loop simulation and closed-loop Cyberpatientin-the-loop experiments for TNF-*α* control. Each line shows the mean of two biological replicates, with shaded regions representing ±SD. (D) Closed-loop performance prediction. Left: Correlation between predicted (by nullgram) and measured (by Cyberpatient-in-the-loop) steady-state cytokine levels (dashed line: *y* = *x*). Right: Relative sensitivity of the different setpoint controllers.

We selected a TNF-*α*–driven rheumatoid arthritis model (*48*), where the therapeutic effector adalimumab represses TNF-*α*. To interface the disease model with living cells, we coupled the LightON optogenetic sensor (*49*) (Supplementary Fig. S3A) to the controller, such that light induction increases activator levels. This implementation was realized using a two-plasmid poly-integration strategy (Supplementary Fig. S3B). One plasmid encoded the LightON system together with a GFP reporter serving as a copy-number proxy for the integrated plasmid. The same construct also encoded an activator-regulated RFP as the controller output, fused to a destabilization domain ecDHFR (G67S/Y100I) (*50*). The other plasmid expressed the splicer IntN and a FRFP marker. The ratio of the FRFP and GFP markers set the controller setpoint.

We used an in-house–developed programmable light-illumination array (*51*) to profile controller dose–response curves across monoclonal controller lines with varying genetic setpoints. As expected, increasing the setpoint shifted the ultrasensitive region of the controller dose–response curve (Supplementary Fig. S3C-D). However, this analysis also revealed a trade-off: higher setpoints were associated with a reduction in maximum controller output expression and a flattening of the response slope. These findings suggest that controllers operating at high setpoints may be constrained by reduced actuation capacity, potentially compromising their ability to reject perturbations.

To interface with the cyberpatient, we selected three monoclonal controller cells spanning low, medium, and high setpoints and profiled their dose–response curves via live-cell microscopy (Supplementary Fig. S4). These measurements showed similar slope flattening at high setpoints as observed in flow-based assays (Fig. 3B, Supplementary Fig. S3D). In parallel, we used the disease model to simulate the drug response curve, which characterizes how TNF-*α* levels respond to changes in adalimumab infusion rate.

As the disease model is defined in terms of TNF-*α* concentration and adalimumab infusion rate, while the controller operates in light–fluorescence space, direct coupling is not possible. Nullgram profiling therefore provided a principled calibration framework to align the input–output relationships of the engineered cells with the disease model (Fig. 3B). Specifically, controller output expression, measured by fluorescence, was mapped to adalimumab production rate, while TNF-*α* was converted into optogenetic light inputs. Both mappings were tunable *in silico*, allowing the intersection between the controller and process curves to be positioned within a desired operating regime. Importantly, these adjustments correspond to biologically meaningful parameters, such as promoter strength, receptor sensitivity, and clonal expression level, that can be modulated through genetic design or cell selection.

Using the calibrated controller–process interface, we leveraged the Cyberpatient-in-the-loop platform (Supplementary Fig. S5) to evaluate closed-loop inflammatory control. Controllers with varying setpoints had comparable growth rates during the experiment (Supplementary Fig. S6B). In contrast to open-loop simulations, closed-loop experiments showed that controllers stabilized TNF-*α* within a physiological range and robustly rejected inflammatory perturbations across patient variability (Fig. 3C). Additionally, closed-loop systems exhibited faster settling and lower steadystate error compared with constitutive (open-loop) control matched to the same steady-state TNF-*α* level (Supplementary Fig. S6C-D). Steady-state TNF-*α* levels (Fig. 3D, left) and adalimumab production rates (Supplementary Fig. S6E) closely matched predictions from nullgram profiling. Notably, controllers operating at higher setpoints exhibited diminished closed-loop performance, validating nullgram-based predictions of reduced sensitivity and actuation capacity (Fig. 3D, right).

Together, these results demonstrate that Cyberpatient-in-the-loop provides a physiologically relevant framework for testing synthetic biological controllers in complex disease environments. By combining nullgram-guided design, real-time cellular interfacing, and computational models, the platform enables rational calibration, scalable validation and predictive evaluation of dynamic therapeutic control systems.

## 3 Nullgram-guided diagnosis reveals limits of TNF-***α*** sensing controllers and guides topology redesign

Analysis of the disease-coupled system using the Cyberpatient-in-the-loop platform together with nullgram profiling revealed consistent underperformance of the controller at high setpoints, coinciding with a pronounced flattening of the controller dose–response curve slope (Fig. 3B). This failure mode is qualitatively distinct from the earlier nullgram profiling performed on a simplified closed-loop system, which attributed high-setpoint limitations primarily to operating-point saturation rather than a loss of controller sensitivity (Fig. 2C). High-setpoint breakdown of the TNF-*α* controller observed in co-culture assays (Supplementary Fig. S1) raised the question of whether this failure results from operating-point saturation alone or from a fundamental loss of controller sensitivity.

To elucidate the mechanism underlying this behavior, we constructed TNF-*α*–sensing controller dose–response curves by placing the activator under an NF-*κ*B–responsive promoter, such that sensing nonlinearity is explicitly embedded in the controller input–output relationship (Fig. 4A). In this setting, the effective controller input is jointly determined by the TNF-*α* level and the integrated copy number of the NF-*κ*B sensor. As a result, the controller setpoint emerges from the ratio between the reference and sensor inputs, similar to the optogenetic controllers used in microscopy-based experiments (Supplementary Fig. S3B), rather than from a single input variable. This analysis revealed a marked reduction in controller dose–response slope at high setpoints (Fig. 4B), recapitulating the failure observed when using the optogenetic sensor (Fig. 3B). We hypothesized that this reduction arises from two interacting effects: nonlinear NF-κB sensing, which may impose an effective ceiling on activator expression, and accumulation of splicing-derived products that retain DNA-binding capacity but lack full activation domains. Such truncated species have been shown to act as dominant-negative repressors, thereby limiting transcriptional output (*52*).

**Figure 4:**
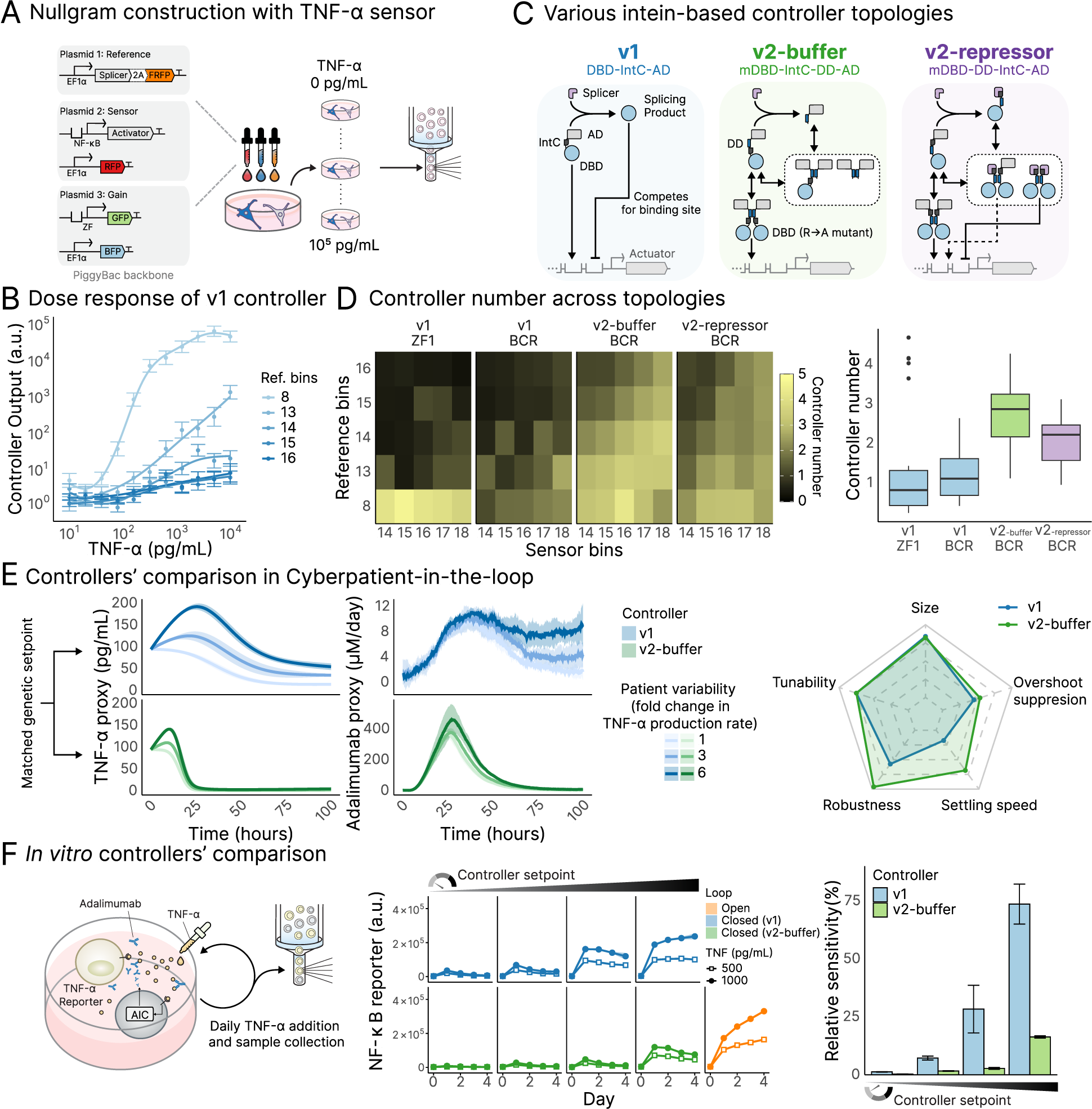
Multi-dimensional nullgram profiling identifies failure mechanisms and guides topology redesign for robust TNF-*α* regulation. (A) TNF-*α* sensing nullgram circuit. The activator is placed under an NF-κB promoter with mRuby3 as copy number proxy, IntN is co-expressed with miRFP670 (P2A, EF1α promoter), and the actuator is mGreenLantern with mTagBFP2 as proxy. A polyclonal library generated by PiggyBac integration was exposed to graded levels of TNF-*α*. (B) Nullgram shows slope reduction at high TNF-*α*. Points are binned single-cell means ± SEM by reference bin; smoothed GAM curves illustrate loss of sensitivity. (C) Various intein-based controller topologies. In the v1 monomeric architecture, IntC is inserted between the activation domain (AD) and the DNA-binding domain (DBD), such that intein-mediated splicing generates DNA-binding splice products. In the v2 dimeric architecture, a mutated DNA-binding domain (mDBD) binds DNA only in the dimeric state. The v2-buffer architecture prevents formation of DNA-binding splice products by placing the intein between the DNA-binding and dimerization domains (DD), whereas the repressor variant produces partially active splice products. (D) Controller numbers across topologies. The controller number is defined as the slope of the log–log relation between actuator output and TNF-*α* input. Buffer maintained high values across setpoints, whereas monomers flattened and the repressor was intermediate. (E) Cyberpatient-in-the-loop comparison of controllers with matched genetic setpoints. Left: Time-courses of TNF-*α* and adalimumab rate proxies. Each line shows the mean of three biological replicates, with shaded regions representing ± SD. Right: Normalized performance scores (0–1 scale) across five metrics: circuit size, tunability, robustness, settling speed, and TNF-*α* overshoot suppression. Scores normalized to fixed caps; larger area indicates better overall performance. (F) Co-culture validation. Both designs suppressed TNF-*α* at low setpoints, but only buffer maintained control at high setpoints. Open squares, 500 pg/mL TNF-*α*; filled circles, 1000 pg/mL. Blue, v1; green, v2-buffer; orange, open-loop. Lines are means of two replicates ± SEM. Bar plot shows relative sensitivity vs open-loop at day 4.

Based on this insight, we sought to eliminate DNA-binding splice products through circuit topology design. In monomeric transcription factor designs, the most feasible site for intein insertion lies between the activation domain (AD) and the DNA-binding domain (DBD), which yields DNA-binding splice products. Although insertion within structured domains could expand the design space, it would require extensive protein engineering to maintain domain function and efficient splicing. In contrast, an alternative strategy to expand the design space is the use of dimeric transcription factors, which introduce additional inter-domain intein insertion sites through the presence of a dimerization domain (DD) (*26*).

We therefore adopted a dimeric transcription factor scaffold based on a BCR zinc-finger with the R39A mutation (*53*), which requires dimerization for DNA binding. We refer to this architecture as version 2 (v2), in contrast to the previously used monomeric designs (version 1, v1). For v2, two distinct topologies were defined by the intein insertion site. In the v2-buffer topology, the intein was placed between the DBD and DD, preventing the formation of DNA-binding splice intermediates. In contrast, in the v2-repressor topology, insertion between the DD and AD still permitted partial transcriptional activation by splice products (Fig. 4C).

To ensure comparable steady-state measurements, all four controller architectures (v1-ZF, v1-BCR, v2-buffer BCR, and v2-repressor BCR) were generated by poly-integration and induced by TNF-*α*, and population-level expression was monitored daily. Most conditions stabilized by day 2; only v1-ZF at high TNF-*α* showed later loss of high-expression cells, so subsequent analyses used day 2 data (Supplementary Fig. S7A). Given the distinct molecular mechanisms underlying each controller topology, we estimated controller dose–response curves using a model-free generalized additive model (GAM) smoothing approach, rather than fitting topology-specific mechanistic models (Supplementary Fig. S7B). The *controller number* was then defined as the maximum local slope of the dose–response curve. These results indicate that v2-buffer maintained high controller numbers across a broad range of setpoints. In contrast, v1 controllers, including ZF1 and a wildtype BCR ZF variant introduced as a DBD-matched control, showed steep dose–response slopes only at low setpoints, followed by marked flattening; the repressor topology exhibited intermediate performance (Fig. 4D).

Based on our previous work (*26*) and the theoretical analysis presented in the Supplementary Note, it can be shown that the v2-buffer controller also exhibits a PID structure, albeit with a different molecular implementation than the v1 controller. While the splice product in the v2-buffer controller does not act as a transcriptional repressor, it buffers activator dimerization (Fig. 4C), thereby regulating the effective pool of active dimers and shaping controller dynamics.

To test the dynamic performance of v2-buffer controller in disease environments, we used the Cyberpatient-in-the-loop platform to compare v1 and v2-buffer controllers, focusing on v1-ZF as the representative monomeric baseline used throughout earlier analyses. The optogenetic implementations of the v1 and v2 controllers exhibited controller numbers consistent with those observed for TNF-*α* sensing (Supplementary Fig. S8A-B). We conducted their comparison at high genetic setpoints, where v1 controllers had previously been shown to fail (Supplementary Fig. S3B, S8C). As predicted, the v1 controller collapsed under these conditions, whereas the v2-buffer controller preserved stable regulation (Fig 4E, left). Notably, the buffer topology maintained effective actuation capacity at high setpoints, enabling sufficient therapeutic output to counteract disease-driven perturbations (Fig 4E, middle). Despite a modest increase in controller size due to the added dimerization domain, the v2-buffer topology consistently outperformed v1 across both static and dynamic performance metrics, including robustness, settling speed, and overshoot suppression (Fig. 4E, right).

Guided by nullgram predictions and supported by Cyberpatient-in-the-loop validation, we next compared v1 and v2-buffer controllers across different setpoints in a co-culture time-course assay. For each topology, four matched setpoint levels were selected to span comparable expression ranges, and responses were monitored daily (Supplementary Fig. S9). Both designs effectively suppressed TNF-*α* at low setpoints, but only the v2-buffer topology maintained control at high setpoints (Fig. 4F). These results highlight that systematically examining controller input–output behavior in highthroughput can efficiently uncover failure modes and guide circuit engineering, thereby streamlining the path from mechanistic insight to functional validation.

## 4 An improved controller enables robust TNF-***α*** regulation in primary human immune cell co-culture

Finally, we evaluated the v2-buffer topology as an application-ready design. In co-culture, the controllers across different setpoints were capable of regulating TNF-*α* levels despite external disturbances (Fig. 5A, left). TNF-*α* reporter signals indicated that the controllers spanned an approximately eightfold dynamic range and scaled with controller setpoints (Fig. 5A, right). These results confirm both the stability and tunability of the v2-buffer controller. To further assess translational relevance, we co-cultured the v2-buffer controller with human peripheral blood mononuclear cells (PBMCs). Varying amounts of T cell activation beads were introduced to simulate different levels of inflammation as external disturbances, and cytokine production was measured 48 hours later. In co-cultures without a controller, TNF-*α* levels scaled with the magnitude of the perturbation. In contrast, the v2-buffer controller effectively suppressed TNF-*α* accumulation, demonstrating effective TNF-*α* regulation (Fig. 5B). As a central mediator of the immune response, TNF-*α* initiates a positive feedback loop that amplifies cytokine release through reciprocal activation between immune cells (*54, 55*). By breaking the TNF-*α*–driven inflammatory feedback loop, the controller not only suppressed TNF-*α* but also reduced downstream cytokine release, including IL-1*β*, IL-6, and IFN-*γ*, demonstrating broad immunomodulatory effects from a single-target intervention (Fig. 5B). Together, these results establish the controller’s functional relevance in primary human immune settings and provide a foundation for future adaptive immunotherapies.

**Figure 5:**
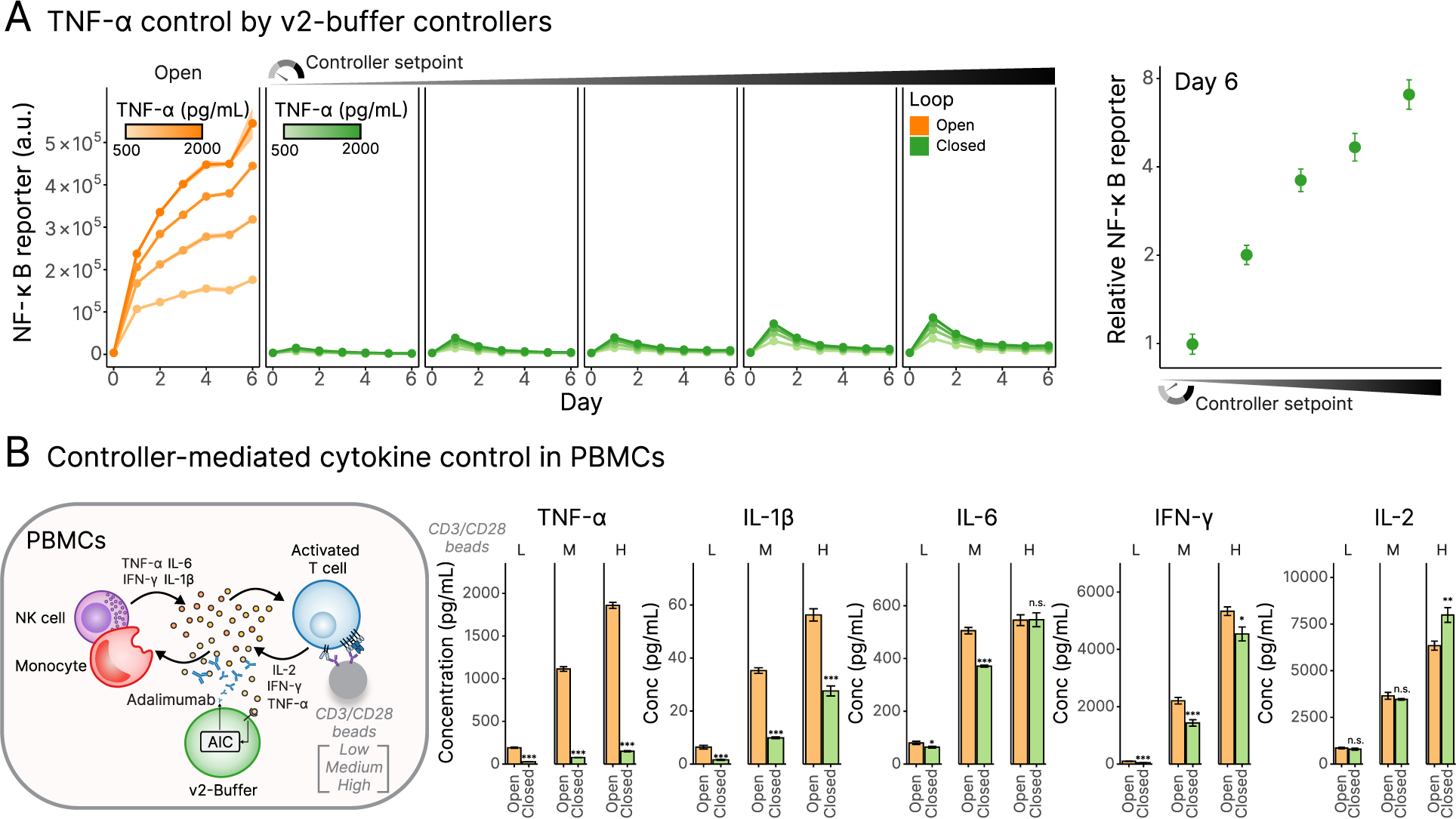
Buffer controller maintains stable setpoints and regulates cytokines in co-culture with primary human cells. (A) Setpoint tracking of the v2-buffer controller in HEK293T coculture. NF-*κ*B reporter output from reporter cells, used as a proxy for extracellular TNF-*α* bioactivity, scaled proportionally with controller setpoints despite TNF-*α* disturbances. Left: Green lines indicate closed-loop buffer AIC and orange lines indicate open-loop controls. Line intensity denotes TNF-*α* concentrations (500, 1000, 1500, and 2000 pg/mL). Each line shows the mean of two biological replicates with shaded regions representing ± SEM. Right: Circles indicate mean NF-*κ*B reporter levels at day 6, normalized to the lowest setpoint and averaged across all disturbance conditions (± SEM; two biological replicates across four TNF-*α* concentrations). Clones are ordered from low to high controller setpoint. (B) v2-buffer AIC-mediated cytokine regulation in PBMCs. Co-culture with human peripheral blood mononuclear cells (PBMCs) suppressed TNF-*α* accumulation across low, medium, and high levels of T cell activation. In addition to TNF-*α* suppression, the controller reduced downstream cytokines IL-1*β*, IL-6, and IFN-*y*, while leaving IL-2 largely unaffected, consistent with selective disruption of a TNF-*α*–driven inflammatory feedback loop. Data represent mean ± SEM of three biological replicates with duplicate ELISA measurements; significance was determined by two-tailed *t* test (\*\*\**P<* 0.001; \*\**P <* 0.01; \**P <* 0.05; n.s., not significant).

## 5 Discussion

In this work, we developed an integrated approach to design and test closed-loop cell therapies with predictable behavior. The framework combines multi-dimensional nullgram profiling, which captures steady-state controller behavior, with Cyberpatient-in-the-loop, a platform for testing engineered cells in dynamic, disease-relevant environments. Together, these tools allowed us to engineer advanced homeostatic cell therapies and achieve robust and precise immune control.

Nullgram profiling provides an experimentally derived steady-state input–output map of a biological controller, allowing direct prediction of closed-loop equilibria, sensitivity limits, and failure modes across architectures and operating regimes. When combined with poly-integration, this approach enables high-throughput comparison of controller designs, thereby guiding topology selection and tuning prior to full closed-loop implementation. More broadly, nullgram analysis reflects a general design principle in synthetic biology: complex nonlinear systems can be engineered by decomposing steady-state behavior into interpretable equilibrium relationships (*42–44*). Building on this interpretability, nullgram profiling enables an alternative design paradigm that moves beyond purely iterative controller engineering. Rather than tuning a single architecture, libraries of controller variants spanning diverse sensing mechanisms, actuation strategies, and internal topologies can be systematically profiled in a single experiment. The developed methodology naturally extends to larger library sizes, as recent massively parallel genetic screening approaches demonstrate the feasibility of assaying large libraries of multi-kilobase gene circuits in mammalian cells (*56*). Taken together, nullgram profiling enables scalable design, comparison, and selection of feedback controllers prior to extensive closed-loop validation, supporting more systematic engineering of closed-loop cell therapies.

To extend these steady-state insights into dynamic disease contexts, we developed Cyberpatientin-the-loop. This framework embeds candidate engineered cell therapies directly into virtual disease environments, allowing assessment of the closed-loop system’s dynamic behavior. Quantitative systems modeling approaches, including PK/PD models, have been widely used in drug development to assess safety, efficacy and predict clinical outcomes (*57, 58*). In some cases, such models have reached sufficient maturity to support regulatory decision-making, exemplified by the UVA/-PADOVA Type 1 Diabetes simulator, which has been approved by the FDA as a surrogate for animal testing (*59, 60*). Our platform builds on this foundation by coupling such disease models directly to engineered cells, enabling real-time evaluation of closed-loop therapeutic strategies that cannot be assessed using simulations or static assays alone. A growing number of curated disease models, available through public repositories such as BioModels (*61*) and through domain-specific efforts in endocrinology (*60, 62*), rheumatology (*63, 64*), and inflammation (*65*), provides a rich resource for extending Cyberpatient-in-the-loop to additional therapeutic contexts. The developed method mirrors hardware-in-the-loop testing paradigms that are widespread in engineering and have been employed in medical device development (*66, 67*), providing a systematic strategy for iterative testing of adaptive cell-based therapies. In parallel, advances in machine learning are enabling the construction of patient-specific digital twins from clinical and molecular data (*68*). Coupling these personalized disease models to Cyberpatient-in-the-loop could allow engineered controller cells to be tested against individualized physiological settings, opening a path towards patient-specific closed-loop cell therapies.

Extending the remarkable successes of cell therapies, such as those achieved in hematological malignancies, to broader indications demands engineering additional capabilities, including improved specificity, persistence, and function in hostile microenvironments (*14, 15, 69*). As a result, cell therapy development becomes a multi-dimensional optimization problem spanning sensing, actuation, and control, while simultaneously introducing new safety considerations associated with programming novel cellular behaviors (*70*). Despite this complexity, existing development often relies on iterative trial-and-error testing, in which circuit architectures are evaluated primarily through full closed-loop testing in specific experimental contexts. Such approaches can obscure the mechanistic origins of failure and hinder the systematic generalization of design insights across applications. Our approach addresses these challenges by enabling high-throughput, predictive iteration grounded in mechanistic insight. Using TNF-*α* regulation as a case study, it enabled the identification of mechanistic limitations arising from sensing nonlinearity and splicing-induced repression and directly motivated the engineering of an advanced topology, which improved therapeutic performance. More broadly, this work highlights how principled, systematic exploration of design space can facilitate the translation of increasingly complex engineered cell behaviors into safer and more predictable therapies.

## Acknowledgments

We thank Dr. Timothy Frei, Dr. Sant Kumar, and Dr. Stanislav Anastassov for their support during the early stage of the project. We thank Dr. Stephanie Aoki, Tomas Kündig, Dr. Filip Bochner, Orlane Maxit, and Cristian Peca for their support and valuable feedback. We thank Dr. Aleksandra Gumienny, Dr. Mariangela Di Tacchio, Dr. Chiara Cavallini, and Dr. Svitlana Malysheva from ETH Zurich D-BSSE Single Cell facility for their support and assistance throughout this study.

## Author contributions

M.K., C.-H.C. and A.A. conceived the project. C.-H.C. and A.A. constructed the circuits and carried out experiments to generate the data. A.A. developed the Cyberpatientin-the-loop platform with assistance from M.C.. M.H. and M.F. performed linear perturbation analysis. F.C. established PBMC isolation and T cell activation protocols. C.-H.C., A.A., and S.B. analyzed the data. M.K., C.-H.C., A.A. wrote the manuscript with input from all authors. M.K. secured funding and supervised the project.

## Data and materials availability

All data are available in the main text or the supplementary materials. Accession codes will be made available prior to publication. Correspondence and request for materials should be addressed to M.K.

## Supplementary materials

Materials and Methods

Supplementary Text

Figs. S1 to S9

Tables S1

References (*7–74*)

